# Simulating isolated populations to identify emerging genetic structure in the absence of selection

**DOI:** 10.1101/853895

**Authors:** C Hosking, R Ogden, H Senn

## Abstract

Conservation efforts are often informed by measures of genetic structure within or between isolated populations. We have established a simulation approach to investigate how isolated or captive populations can display misleading (i.e recently acquired) genetic structure as a result of genetic drift. We utilized a combination of softwares to generate isolated population genetic datasets that allow interrogation of emerging genetic structure under a range of conditions. We have developed a new statistic, S, to describe the extent of differentiation due to genetic drift between two isolated populations within the clustering software, STRUCTURE.

A novel method to infer the effects of genetic drift on structure among isolated populations

**Graphical Abstract:** 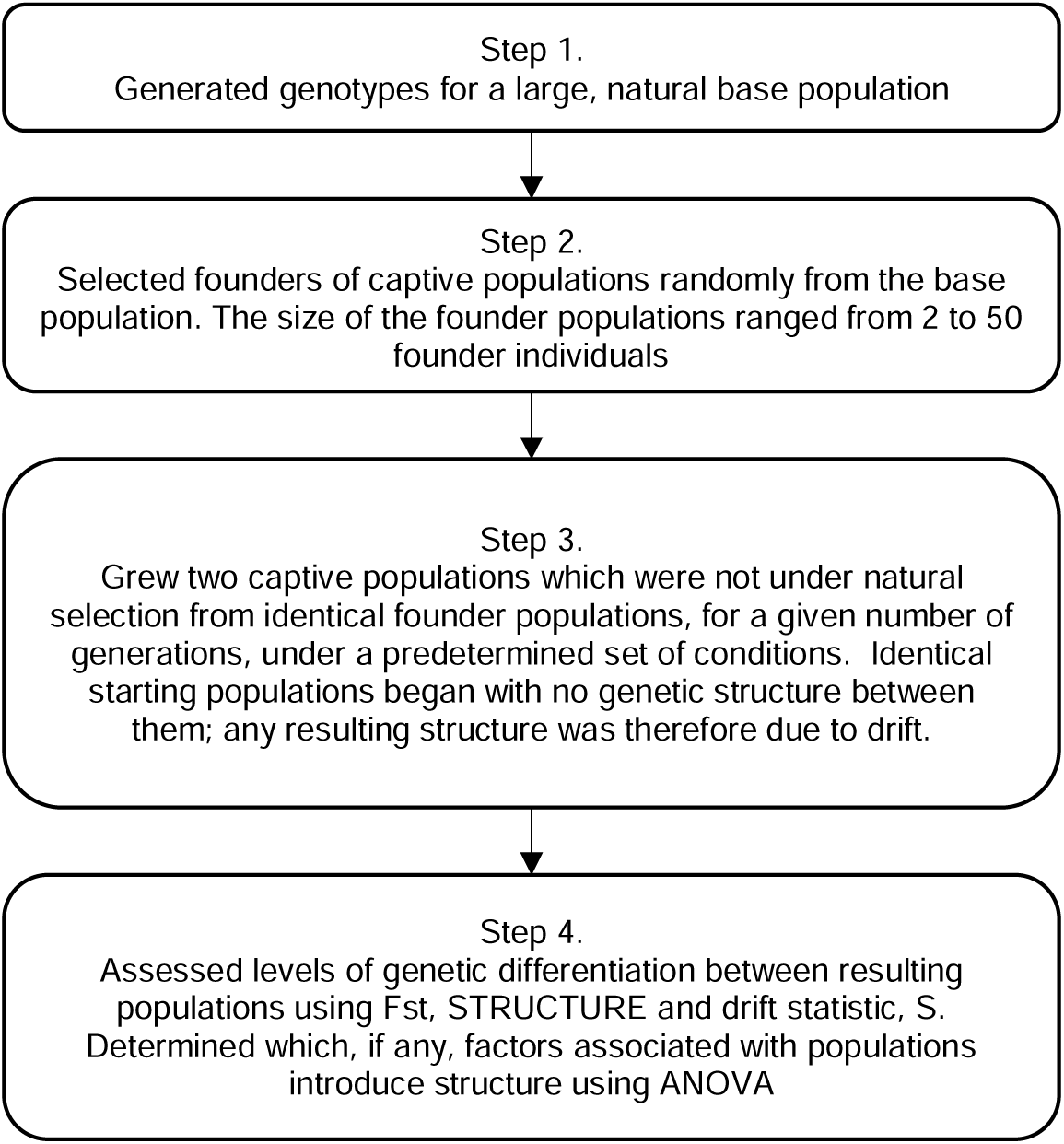

**SPECIFICATIONS TABLE:** 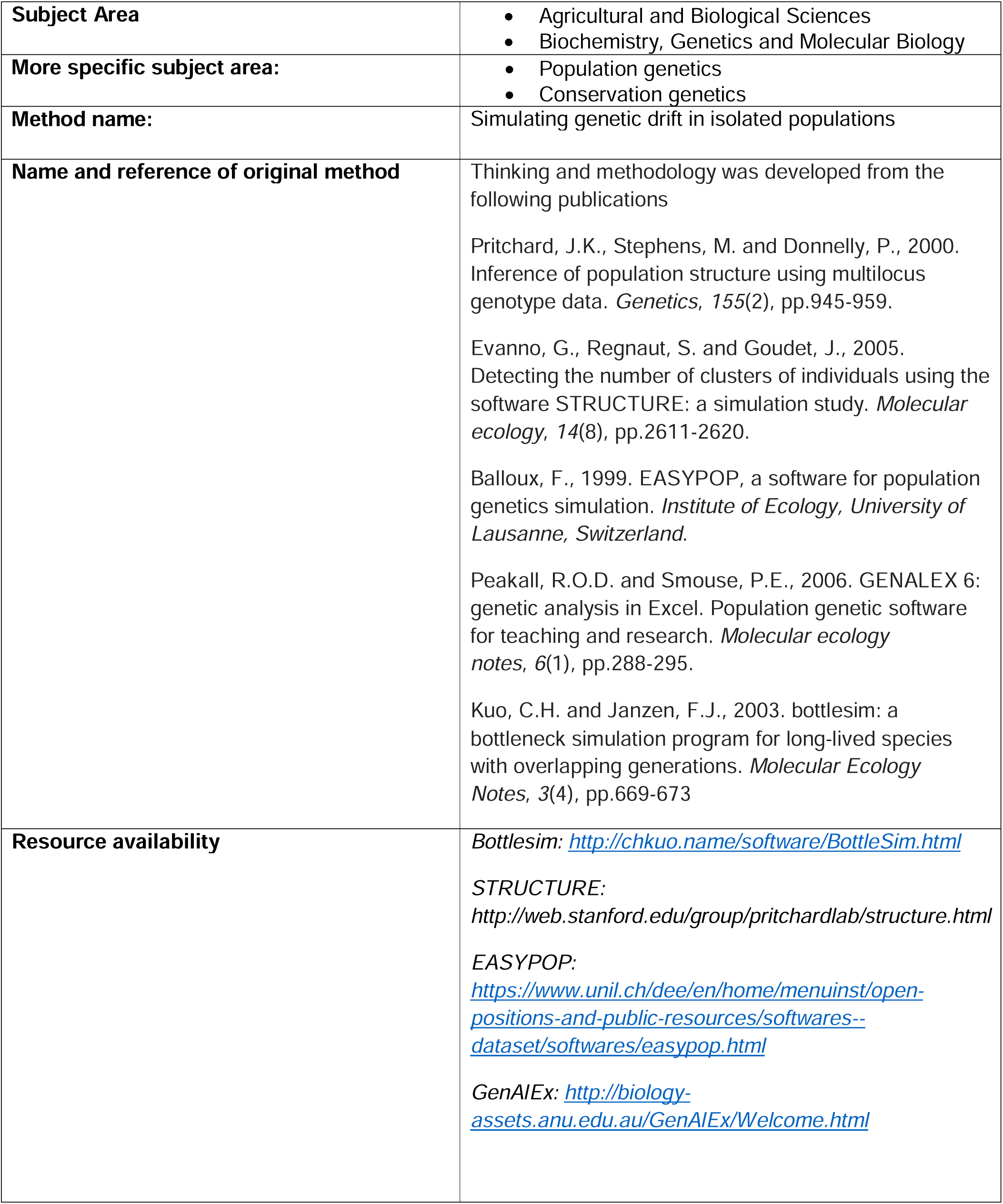

## Method details

Conservation efforts are often informed by the genetics of isolated populations, either remaining in the wild or in captive breeding programmes. Populations such as these are at risk of strong genetic drift from the original wider population and subsequent differentiation may no longer represent meaningful or adaptive genetic variation. To investigate this effect, we have chosen a simulation approach which demonstrates the potential for genetic drift to lead to identifiable genetic structure within a metapopulation. This approach removes background variation which would be present in attempts to address the question using empirical data. Simulations begin with identical starting populations and thus, an initial F_ST_ of zero. Any detected genetic structure present at the end of the simulation must therefore be attributed to genetic drift, rather than any historical evolutionary forces. The simulation approach followed that shown in Figure 1 and is presented in more detail below.

**Figure 1.**
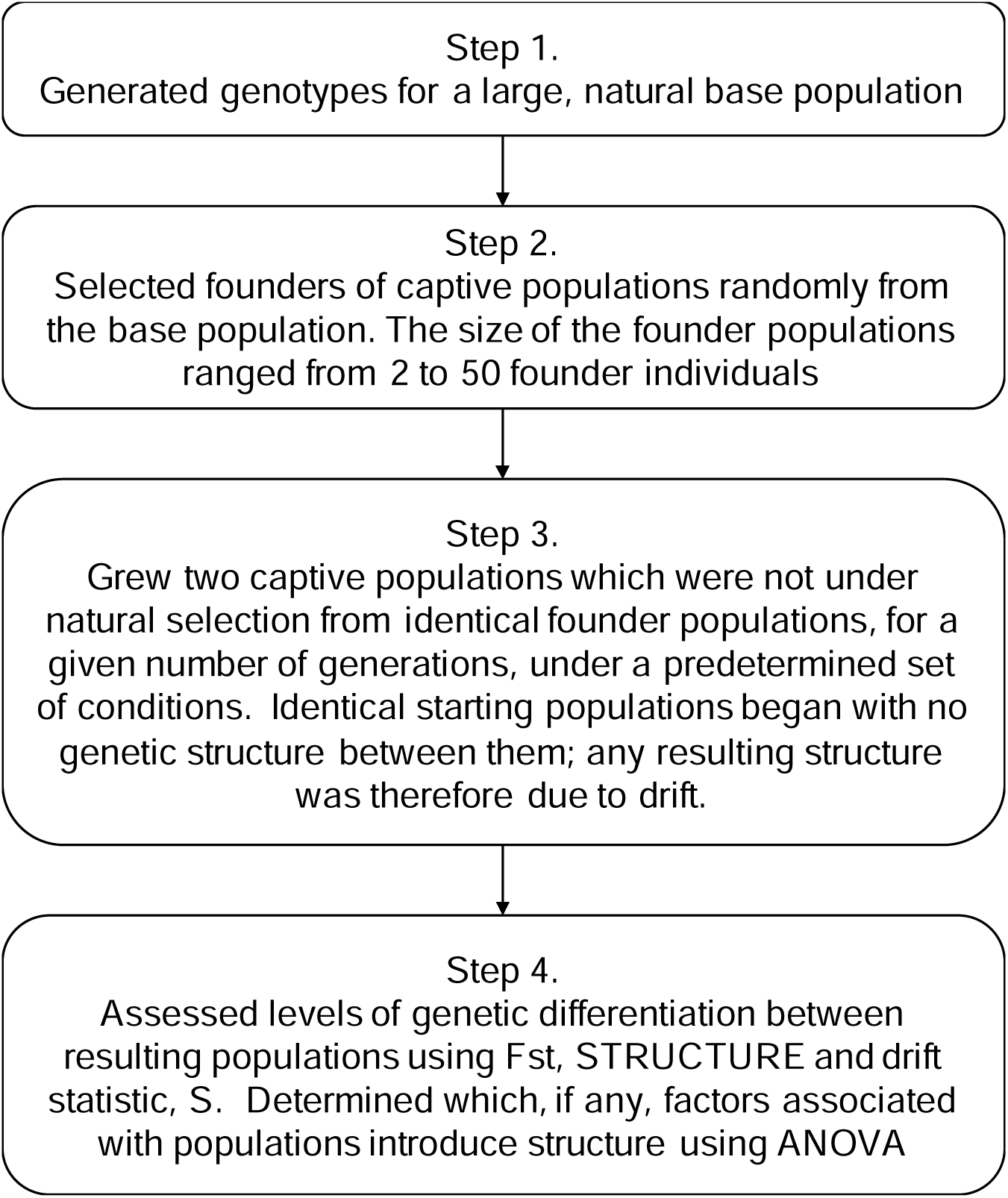
Outline of the simulation process, including the production and growth of isolated populations and their subsequent analysis

### Step 1: Simulating a panmictic population

In order to simulate individuals being removed from wild populations into captive breeding programmes it was necessary to simulate a large, diverse, panmictic population from which to sample (see Figure 1). The parameters for this wild population were based on values describing the red deer population of Scotland. This metapopulation and its structure has been studied for many years (Nussey *et al*. 2006; Perez-Espona *et al*. 2008; Pérez-Espona *et al*. 2013). Although it cannot be considered a truly panmictic population due to evidence of structure primarily as a result of geographical barriers, the population is large, diverse and healthy (Perez-Espona *et al*. 2008; Pérez-Espona *et al*. 2013). The ungulate template is continued throughout the parameters simulated. EasyPop 2.0.1 software was used to simulate the wild population (Balloux 1999), see parameters in Table 1, so that there is little or no obvious structure in the larger population. Simulated microsatellite loci were assumed to be unlinked. Individuals were initially randomly assigned a genotype and the population was simulated for 100 generations. This ensured that the population had not gone to fixation as a result of drift at any loci, but that some alleles had been lost.

**Table 1.** Simulated wild population parameters. 80% of microsatellite mutations were single step mutations (SSM) and 20% were mutations between allelic states at an equal rate (KAM) (Balloux and Lugon-Moulin, 2002). ^†^ indicates value taken from Kruglyak et al. (1998), * indicates values taken from Phillips et al (2008)

| Parameter | Set |
| --- | --- |
| **N** | 1000 |
| **Mating System** | Random |
| **Sex Ratio** | Equal |
| **Microsatellite mutation rate (per locus per generation)** | $5 \times 10^{-4\dagger}$ |
| **Microsatellite mutation model** | SSM and KAM |
| **Maximum number of microsatellite alleles** | 14 |
| **SNP mutation rate (per locus per generation)** | $2.5 \times 10^{-8*}$ |
| **SNP mutation model** | KAM* |
| **Number of SNP alleles** | 2 |

### Step 2: Select founders

The EasyPop simulation results include the genotypes for every resulting individual in the population in a format which can be imported into Microsoft Excel using the GenAlEx v6.5 plugin (Peakall and Smouse 2006). The random number function within Excel was then used to assign a random value between 0 and 1 to each individual. When ranked smallest to largest the top individuals were used as founders for the captive population. For each subsequent simulation with a different founder population size the simulated wild population was randomly sorted again and the individuals selected. The result being that every simulation with a given founder population size began with the same individuals but a new set of randomly selected individuals was chosen for every alternative founder population size investigated.

### Step 3: Grow captive populations

Simulations were designed primarily to test the effects of time in isolation and the number of founders on the rate at which population structure appears due to drift. Additionally, the effects of mating system, population growth rate and the ability of alternative marker numbers and types (microsatellite versus SNPs) to detect population structure were investigated to ensure results were not limited to a narrow set of parameters. Simulated captive populations were produced using BottleSim v 2.6 (Kuo and Janzen 2003) from the genotypes of individuals selected from the wider population as described in Step 2.

All simulations followed a diploid, multilocus model with a variable population size. The population size and sex ratio at every generation was defined prior to simulation. Longevity was set to 15 years (Price 1989) and had a generational overlap of 100%. This maximal generation overlap allowed all individuals who had reached reproductive maturation (see below) to be considered potential parents. Mating takes place every simulated year and generations are overlapping, therefore results are discussed in terms of generations. Reproductive maturation was set to zero in order to prevent simulation failure due to a lack of suitable breeding individuals in such small populations as those represented here.

In each case we compared population differentiation between two simulated captive populations bred from identical starting (sub)populations. As the starting populations are identical, there is no genetic structure between the two populations at time zero. Any structure found between the resulting populations following simulated population growth must therefore be the result of drift (since the model contains no selection). This is an extreme scenario, which is perhaps unrealistic, but conservative. However, if we detect structure here we can assume that real life populations would actually be differentiated even further if they are started from similar but non-identical sources (for example two groups of animals selected from the same wild populations).

#### Founder population size

The number of founders used to represent captive populations covers the range found in the literature for zoo populations (Armstrong *et al*. 2011: *Addax nasomaculatus -* 2; Beauclerc *et al*. 2010: *Peltophryne lemur* - 4 and 38 founders of Northern and Southern populations, respectively; Marsden *et al*. 2013: *Lycaon pictus* - 38; McGreevy *et al*. 2011: *Dendrolagus matschiei* - 19 and Price 1989: *Oryx leucoryx -* 9). The core scenarios were carried out using individuals genotyped at 10 microsatellite loci (this represents a typical number used for conservation genetics), under a constant 10 % population growth rate and with random mating with a limiting carrying capacity of 200 individuals, as is the norm in European captive populations of large ungulates. The following parameters were also investigated: a one male mating per generation mating system, micro satellites versus SNP loci and the ability of various numbers of markers (microsatellites: 5 - 40, SNP: 48 - 384) to detect population structure.

#### Number of generations in isolation

Simulations were run for a maximum of 15 generations. These same starting populations were rerun for 2, 5 and 10 generations of population growth in isolation. The resulting metapopulations were analysed for any evidence of genetic structure (Step 4).

### Step 4: Analysis of resulting metapopulation

BottleSim output includes the genotype for each individual in the final generation of the simulation. This can be imported into Excel and analysed using GenAlEx v 6.5 (Peakall and Smouse 2006). F_ST_ was calculated for the metapopulation comprised of two subpopulations with identical origins, using Equation 1 from (Nei 1977).

The estimation of F_ST_ within GenAlEx when applied to this kind of data set uses Nei’s equivalent approach adapted for multiallelic loci such as microsatellites, sometimes termed G_ST_, and is calculated as shown,

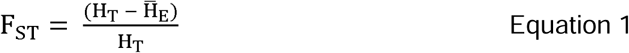

where H_T_ refers to the total expected heterozygosity and H_E_ is the mean expected heterozygosity across populations (Nei 1977).

STRUCTURE v2.3.3 (Pritchard *et al*. 2000) was used to detect any evidence of population structure by clustering individuals based on allele frequencies under the assumptions that clusters are in Hardy-Weinberg Equilibrium and linkage equilibrium. Models for K = 1 - 3 were tested, to ensure differentiation did not continue further than expected, following a burn–in period of 1 × 10^5^ steps and 2 × 10^5^ subsequent MCMC iterations. STRUCTURE modelling of each value of K was repeated three times. The admixture model and correlated allele frequencies among populations were assumed. No prior information regarding population origin was included. The K number of clusters was determined using the method recommended by (Pritchard *et al*. 2000) to maximise the negative likelihood for the model for each value of K. The Evanno *et al*. (2005) method, although popular (Kraus *et al*. 2013; Marsden *et al*. 2012; Nsubuga *et al*. 2010; Row *et al*. 2012; Witzenberger and Hochkirch 2013) is inappropriate for use here as the calculation of the delta K statistic uses likelihood for each sequential value of K and can therefore not be used to identify a panmictic population where K = 1. As a result, all simulated metapopulations were found to have K = 2 using the Evanno method, as did the initial starting populations prior to any population growth. In order to quantify the extent of differentiation between the resulting populations a new structure differentiation statistic, S, has been developed based on STRUCTURE output in the case where K=2, which we have assumed to be the case here.

For each value of K, STRUCTURE calculates Q values for each individual. Q values refer to the proportion of each individual which has been assigned to each of K clusters. Q values are represented in a STRUCTURE bar plot. For each individual when K = 2, there are two values of Q: Q_1_ is the proportion of an individual which can be assigned to cluster 1, Q_2_ is therefore the remaining proportion indicating assignment to cluster 2. These values sum to one. In this study STRUCTURE is being used to detect population differentiation between two populations. The results of a STRUCTURE run can be summarised using the average of these numbers for each individual as shown in Table 2, where

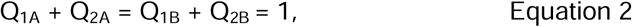

and

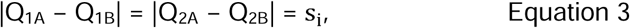

**Table 2.**
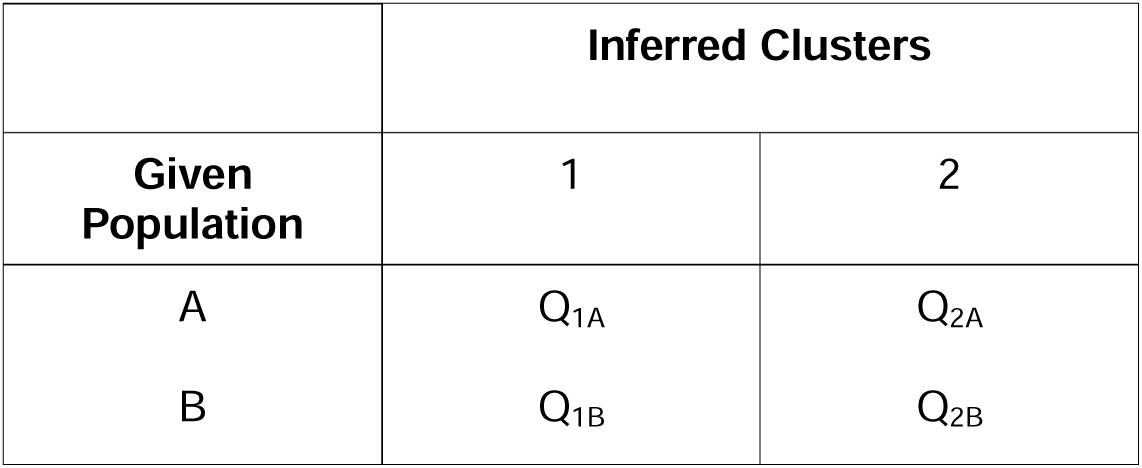
Average proportion of membership of each individual from pre-defined populations to each of the inferred clusters when K = 2. As each value describes an average proportion their values are constrained 0 ≤ x ≤ 1.

| Given Population | Inferred Clusters |  |
| --- | --- | --- |
|  | 1 | 2 |
| A | $Q_{1A}$ | $Q_{2A}$ |
| B | $Q_{1B}$ | $Q_{2B}$ |

Where i refers to the subsequent repetitions, 1 to *n*, of the K = 2 STRUCTURE model. It should be noted that Q_1A_ and Q_2A_ are not constrained by the values of Q_1B_ and Q_2B_. From Table 2 and Equation 3, S is calculated thus,

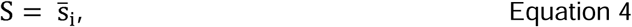

where 0 ≤ S ≤ 1. When S = 0, each individual has been equally assigned to both inferred clusters, suggesting that there is no differentiation between clusters. This can be seen clearly in a STRUCTURE bar plot (Figure 2a) and recommended by (Pritchard *et al*. 2000) as visual confirmation of panmixia. When S = 1, this suggests complete differentiation between clusters. As a result, individuals from each of the original populations have been completely assigned to a corresponding cluster (Figure 2b). However, the major advantage of a quantitative description of STRUCTURE output is its application when visual outputs are hard to interpret, as in Figure 2c. A corresponding value of S was calculated for each simulation in addition to mean F_ST_.

**Figure 2.**
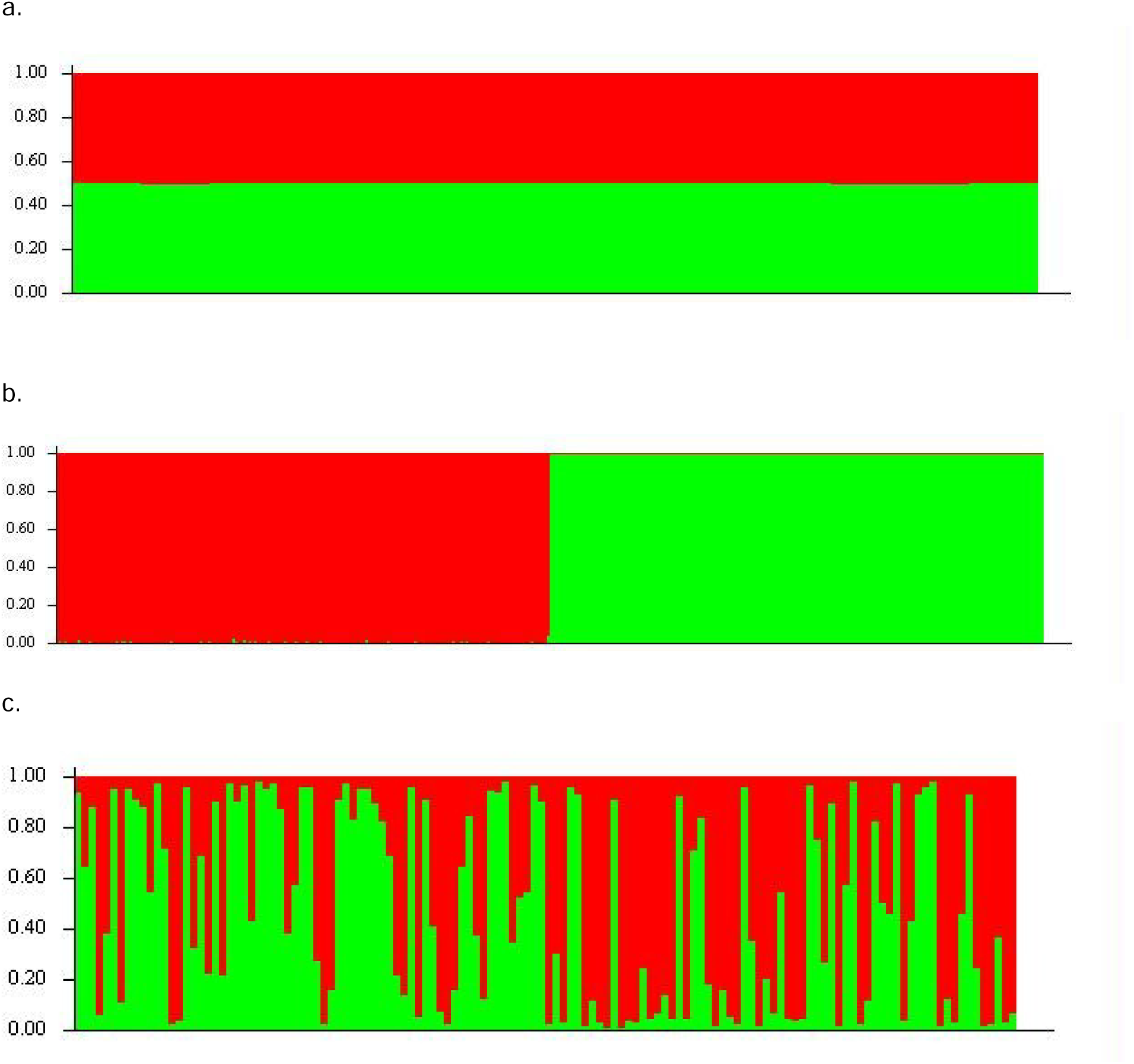
STRUCTURE output from a selection of possible values of S. Each individual is represented by a single vertical bar and colours represent inferred cluster, a) S = 0.0003, b) S = 0.9917 and c) S = 0.2877 where there is evidence to suggest an undifferentiated population as S < 0.5

In summary, we have established a workflow using freely available software to simulate isolated, captive populations and interrogate the appearance of genetic structure as a result of genetic drift alone. Although we have used ungulates and more specifically, red deer, as a model organism to establish the parameters of our simulations, this approach could be used to inform the interpretation of genetic structure in other species of interest.

## Supplementary material *and/or* Additional information

The methodology was developed during supervision of a MSc project thesis in collaboration with the University of Edinburgh’s Institute of Evolutionary Biology: Hosking C. (2013). *Genetic drift may hinder identification of genetic structure in captive populations of endangered species.* Unpublished master’s thesis, School of Biological Sciences, University of Edinburgh.

